# Kv3 channels contribute to the excitability of sub-populations of spinal cord neurons in lamina VII

**DOI:** 10.1101/2021.11.29.470360

**Authors:** Pierce Mullen, Nadia Pilati, Charles H. Large, Jim Deuchars, Susan Deuchars

## Abstract

Autonomic parasympathetic preganglionic neurons (PGN) drive contraction of the bladder during micturition but remain quiescent during bladder filling. This quiescence is postulated to be due to recurrent inhibition of PGN by fast-firing adjoining interneurons. Here, we defined four distinct neuronal types within lamina VII of the lumbosacral spinal cord, where PGN are situated, by combining whole cell patch clamp recordings with k-means clustering of a range of electrophysiological parameters. Additional morphological analysis separated these neuronal classes into parasympathetic preganglionic populations (PGN) and a fast firing interneuronal population. Kv3 channels are voltage-gated potassium channels (Kv) that allow fast and precise firing of neurons. We found that blockade of Kv3 channels by tetraethylammonium (TEA) reduced neuronal firing frequency and isolated high-voltage-activated Kv currents in the fast-firing population but had no effect in PGN populations. Furthermore, Kv3 blockade potentiated the local and descending inhibitory inputs to PGN indicating that Kv3-expressing inhibitory neurons are synaptically connected to PGN. Taken together, our data reveal that Kv3 channels are crucial for fast and regulated neuronal output of a defined population that may be involved in intrinsic spinal bladder circuits that underpin recurrent inhibition of PGN.

**Significance statement:** Neural circuits in the spinal cord and pons mediate the micturition reflex. The spinal cord drives bladder contraction during micturition through the activation of parasympathetic preganglionic neurons in lamina VII of the lumbosacral spinal cord. Despite the significant contribution of these neurons to a crucial physiological reflex, neurons in this region have been under-characterised. This study therefore elucidated and thoroughly characterised distinct neuronal populations in this lamina; we propose that these populations included a fast firing interneuron and subtypes of parasympathetic preganglionic neurons. Further investigation revealed the critical importance of Kv3 channels in the fast firing ability of the interneurons, as well as in synaptic release onto parasympathetic preganglionic neurons.

## Introduction

Autonomic parasympathetic preganglionic neurons (PGN) are situated at the intersection between the lumbar and sacral spinal cord in the intermediolateral laminae, and drive the contraction of the bladder detrusor muscle during the micturition reflex (de Groat and Ryall, 1968a; Morgan et al., 1979; Fowler et al., 2008; de Groat et al., 2014). The quiescence of PGN during bladder filling has been postulated to be due to recurrent inhibition mediated by parasympathetic axon collaterals and fast firing interneurons in the vicinity of PGN (de Groat and Ryall, 1968b; Shefchyk, 2001). A description of neurons in this area and the mechanisms that confer the fast-firing phenotype is lacking, but one strong possibility for a biophysical mechanism promoting fast firing is the expression of Kv3 channels.

Kv3 voltage-gated potassium channels play an important role in shaping a fast firing phenotype and precise synaptic output of neurons (Rudy and Mcbain, 2001). This role is conferred by two important properties; channel activation at relatively high voltages only achieved during an action potential, and fast activation and deactivation kinetics that ensure a rapid repolarisation of the membrane and a short refractory period. Rapid repolarisation produces brief action potentials (APs) which limit calcium influx and thus neurotransmitter release at the pre-synaptic terminal, whereas short refractory periods, in addition to brief APs, allow the neuron to fire APs in quick succession (Kaczmarek and Zhang, 2020). These channels, therefore, are crucial in determining the ability of a neuron to produce a fast but precise output.

Kv3 channels are typically expressed in the soma, nodes of Ranvier and synaptic terminals of neurons in various regions of the central nervous system (CNS); from the auditory brainstem to the cortex (Weiser et al., 1995). Expression has also been observed in the thoracic spinal cord, particularly in dorsal horn interneurons, Renshaw cells and intermediolateral laminae typically associated with sympathetic preganglionic neurons (Deuchars et al., 2001; Brooke et al., 2002, 2006; Song et al., 2006; Nowak et al., 2011).

However, the connection between parasympathetic preganglionic output from the spinal cord and Kv3-mediated fast firing of neurons in the vicinity of PGN hasn’t been established. We thus hypothesised that Kv3 channels are indeed expressed in putative interneurons in the autonomic intermediolateral lamina of the lumbosacral sacral spinal cord and are responsible for facilitating a fast firing phenotype, making them ideal candidates for autonomic inhibition.

## Methods

### Spinal cord tissue preparation

C57bl6 mice (p10-21), according with the UK animals (Scientific Procedures) Act 1986 and ethical standards set out by the University of Leeds Ethical Review Committee, were anaesthetised by intraperitoneal (i.p) injection of sodium pentobarbitone (Euthanal, 60 mg/kg). Upon complete loss of pedal withdrawal, a transverse laparotomy was carried out to remove the ventral ribs and expose the heart. The right atrium was incised and 20 ml of artificial cerebrospinal fluid with high sucrose (sucrose aCSF; sucrose, 217 mM; NaHCO3, 2 mM; KCl, 3 mM; MgSO4.7H2O, 2 mM; NaH2PO4, 2.5 mM; glucose, 10 mM; CaCl2, 1 mM) oxygenated (95% O2/5%CO2) on ice was perfused through the left ventricle and into the circulation system. The mouse was decapitated, the skin removed, and a dorsal laminectomy carried out to expose the spinal cord which was removed following cutting of the rootlets attached to the cord.

Upon removal, the spinal cord was placed in a petri-dish containing ice-cold sucrose aCSF under a dissecting microscope (SM2 2B, Nikon) and the meninges that ensheathe the cord were removed with fine forceps. Lumbo-sacral segments of the spinal cord were set in 3 % agar in aCSF (NaCl, 124 mM; NaHCO3, 26 mM; KCl, 3 mM; MgSO4.7H2O, 2 mM; NaH2PO4, 2.5 mM; Glucose, 10 mM; CaCl2, 2 mM), mounted against a 4 % block of agar for stability using superglue and sectioned in a bath of oxygenated ice-cold sucrose aCSF using an Integraslice 7550 PSDS (Campden Instruments, UK) microtome. Transverse sections were cut at 250-400 μm and incubated in an oxygenated holding chamber containing aCSF and allowed to recover for an hour before recording.

### Patch clamp of spinal neurons

Recordings were carried out at 34 °C using an inline solution heater and temperature controller (SH27B, TC-344C, Warner Instruments, USA). Slices were transferred to and immobilised in an incubation chamber perfused with oxygenated aCSF at a rate of 3-5 ml/minute. Thick walled borosilicate glass microelectrodes (inner diameter 0.86 mm, outer diameter 1.4 mm) were fabricated using a Sutter P97 micropipette puller (Sutter Instruments, USA) with resistances of 5-9 MΩ. The recording patch and bath electrodes used a silver/silver chloride (Ag/AgCl2) interface. Patch microelectrodes were filled with an intracellular solution composed of; K gluconate, 110 mM; EGTA, 11 mM; MgCl2, 2mM; CaCl2, 0.1 mM; HEPES, 10 mM; Na2ATP, 2 mM; Na2GTP, 0.3 mM and 0.5% Neurobiotin, pH 7.4, 290 mOsm (Vector Laboratories, USA). Recordings were obtained at 50 kHz, filtered through a Bessel low pass filter at 10 kHz using a MultiClamp 700A (Axon Instruments, USA), digitised using a Digidata 1322A (Axon Instruments, USA) and recorded in pClamp9 software. An upright microscope (Olympus BX50W1), camera (QImaging Rolera-XR, QImaging, Canada) and QCapture software (QImaging, Canada) were used to visualise the spinal cord section and centre the stage over the region of interest e.g. the lateral region of the lumbo-sacral spinal cord sections.

In current clamp configuration, neurons were characterised by long (1 second) hyperpolarising and depolarising current steps from a holding membrane potential of -70 mV. The holding membrane potential was corrected for a liquid junction potential of -15 mV. Passive properties such as cell capacitance and resistance were monitored with a 50 pA hyperpolarising pulse at the end of each sweep by fitting an exponential function to the voltage decay. Neuronal firing frequency was calculated during 1 second current injections, incrementally increasing by 10 pA. The following frequency of neuronal firing was assessed by applying trains of 10 ms square current pulses to neurons at 20 Hz, 50 Hz and 100 Hz at increasing current amplitudes. The rate of failure (%) to evoke an action potential at each frequency and current pulse was quantified.

In voltage clamp configuration, outward plateau and tail currents were measured during 250 ms, 10 mV incrementing steps from -56 mV. All voltages were corrected for the liquid junction potential and series resistance and holding currents were < 30 MΩ and < 300 pA. Recordings where series resistance changed by >20% were excluded. Non-inactivating Kv currents were isolated by inactivating channels with a 2 second voltage step to -36 mV before applying test voltage steps.

Post-synaptic currents were evoked by a brief pulse (6-8V) using a bipolar electrode positioned in the lateral white matter where descending tracts are located. Excitatory and inhibitory post-synaptic currents (EPSCs and IPSCs) were recorded by holding the neuron at - 56 mV for EPSCs and at 0 mV for IPSCs. Paired pulse stimulation with an inter-pulse duration of 100 ms was used to assess presynaptic potentiation and attenuation. A 10 Hz train of stimulation was also used to assess the synaptic response to repetitive input.

Recordings in control aCSF were started 4 minutes after achieving the whole cell configuration. TEA and dendrotoxin (DTX) were dissolved in extracellular solution to obtain a bath concentration of 0.5 mM and 10 nM, respectively. Recordings were obtained after 8 minutes of dialysis for DTX experiments and after 4 minutes of dialysis for TEA experiments. Passive properties were monitored throughout and recordings where series resistance changed by >20% were excluded.

### Morphological reconstruction

Recorded neurons were filled with 0.5% Neurobiotin and spinal cord slices were fixed on slides with 4 % paraformaldehyde (PFA) for 2 hours. Fixed sections were then washed, permeabilised and incubated in Steptadavidn-555 solution (1:1000, Phosphate buffer (PB) -0.3% Triton X-100). Sections were imaged using a Zeiss LSM880 Upright confocal microscope at 20x and traced using the SimpleNeuriteTracer plugin in ImageJ. Sholl analysis was performed with the plugin and a custom python script was used to compute the angles between endpoints of neurites and the soma.

### Immunohistochemistry

Female wild-type C57BL/6 (3-month-old) were anesthetized with intraperitoneal pentobarbitone sodium (Sagatal, 60 mg/kg) and perfused transcardially with 4% paraformaldehyde (PFA) in 0.1 M phosphate buffer (PB), pH 7.4. After fixation, spinal levels L1, L6, S1 were dissected and incubated in 0.1M PBS containing 30 % sucrose until the tissue sank to the bottom before being embedded and frozen in Surgipath FSC 22 Clear Frozen Section Compound (Leica) freezing medium on dry ice. 20 μm sections were cut using a Leica CM1850 cryostat cooled to approximately -15 °C and mounted onto Superfrost plus (Menzel-Glaser, Thermo Scientific) slides. Sections were washed 3 times in PBS, incubated in 10 mM sodium citrate at 80°C for 20 minutes for antigen retrieval, washed a further 3 times in PBS and blocked and permeabilised for 1 hour in 5 % goat and donkey serum in PB (0.3 % Triton X-100). All primary antibodies were incubated overnight in PB (0.3% Triton X-100), washed in PBS and incubated for 1 hour for directly conjugated secondary antibodies, for 2 hours for biotinylated secondary antibodies and for 30 minutes for streptavidin to avoid endogenous biotin labelling. Antibodies used are listed in Table 1.

**Table 1.** Antibodies and concentrations used

| Target | Supplier | Species Raised in | Dilution | Secondary Detection | Cat. No | Reference |
| --- | --- | --- | --- | --- | --- | --- |
| Kv3.1b | Neuromabs/Antibodies Inc | Mouse | 1:100 | Biotinylated α-mouse IgG1 (Invitrogen A10519), Streptavidin Alexa 555 (Invitrogen) | 75-041 | (Soares et al., 2017) |
| Kv3.3 | Neuromabs/Antibodies Inc | Mouse | 1:100 | Biotinylated α-mouse IgG1 (Invitrogen A10519), Streptavidin Alexa 555 (Invitrogen) | 75-354 | (Soares et al., 2017) |
| GlyT2 (Glycine Transporter) | Synaptic Systems | Rabbit | 1:2000 | Donkey α-Rabbit Alexa 488 (Life Technologies, A21206) | 272 003 | (Jursky and Nelson, 1995) |
| VGAT (Vesicular GABA Transporter) | Synaptic Systems | Rabbit | 1:2000 | Donkey α-Rabbit Alexa 488 (Life Technologies, A21206) | 131 003 | (Tozuka et al., 2005) |
| VGLUT2 (Vesicular Glutamate Transporter) | Synaptic Systems | Rabbit | 1:2000 | Donkey α-Rabbit Alexa 488 (Life Technologies, A21206) | 135 403 | (Zhu et al., 2018) |
| ChAT (Choline acetyltransferase) | Abcam | Goat | 1:1000 | Donkey α-Goat Alexa 488 (Life Technologies, A21206) | Ab18736 | (Merolli et al., 2019) |

### Confocal and Airyscan Imaging

Images of neurons were acquired with the nucleus in view. DIC was used to identify the cell outline and determine a visible nucleus whilst super-resolution Airyscan image were taken of the neurons with the Alexa 555 and Alexa 488 fluorophores being stimulated separately and emissions collected with appropriate bandpass filters for the fluorophores. At least nine neurons from three sections at L6-S1 were measured per animal (n=3). The neuron perimeter was traced in ImageJ to create a region of interest (ROI). A 3 μm band around the cell was created from 2 μm outside the perimeter to 1 μm inside the perimeter. This formed a ROI from which synaptic immunoreactivity and Kv3 puncta in very close apposition with the cell membrane could be segmented. Synaptic immunoreactivity, referred to as boutons herein, and Kv3 puncta were segmented in ImageJ. For object-based co-localisation, co-localisation was defined as the centre of a Kv3 punctum coinciding with the area of a bouton. This was performed using the JaCOP plugin (Cordelieres et al, 2006) on ImageJ and reported as the percentage of co-localised boutons and as the percentage of co-localised puncta.

### Clustering of neuronal e-types

Neurons were clustered based on seven features of neuron firing. These were frequency of action potentials, adaptation index, regularity of firing (interspike interval coefficient of variation), accommodation, as well as the presence of bursts, pauses or delays in firing. Ninety six neurons from 35 mice were clustered on these features, after normalisation, using SciPy K-means with the ideal number of clusters determined by an elbow plot. Principal component analysis (PCA) was performed using the SciPy package and used reduce the electrophysiological parameter dimensions to visualise the k-mean clusters.

### Experimental Design and Statistical Analysis

Custom Python scripts were used to extract and measure action potentials, synaptic currents and outward currents. All statistics were carried out on GraphPad prism software with single neurons representing the experimental unit. Shapiro-Wilks and Levene’s tested normality and homogeneity of variance assumptions. Kolmogorov-Smirnov test was used statistically compare synaptic amplitude distributions. All statistical tests used are described alongside significant results. Values are given as mean ± SD.

## Results

### Kv3 subunits are expressed in the lumbosacral spinal cord

Immuno-labelling of Kv3 subunits Kv3.1b and Kv3.3 revealed widespread expression in the lumbosacral spinal cord, throughout the dorsal, lateral and ventral gray matter, in agreement with previous descriptions of Kv3 subunits in other spinal segments (Brooke et al., 2002, 2006; Song et al., 2006; Nowak et al., 2011) (Fig 1A, B). Magnified images indicated a punctate expression profile within or closely apposed to the somatic and perisomatic membrane of neurons within the intermedio-lateral autonomic zone (Fig 1B). Here we define this region as within lamina VII, lateral to the central canal and adjacent to the gray and white matter border.

**Figure 1.**
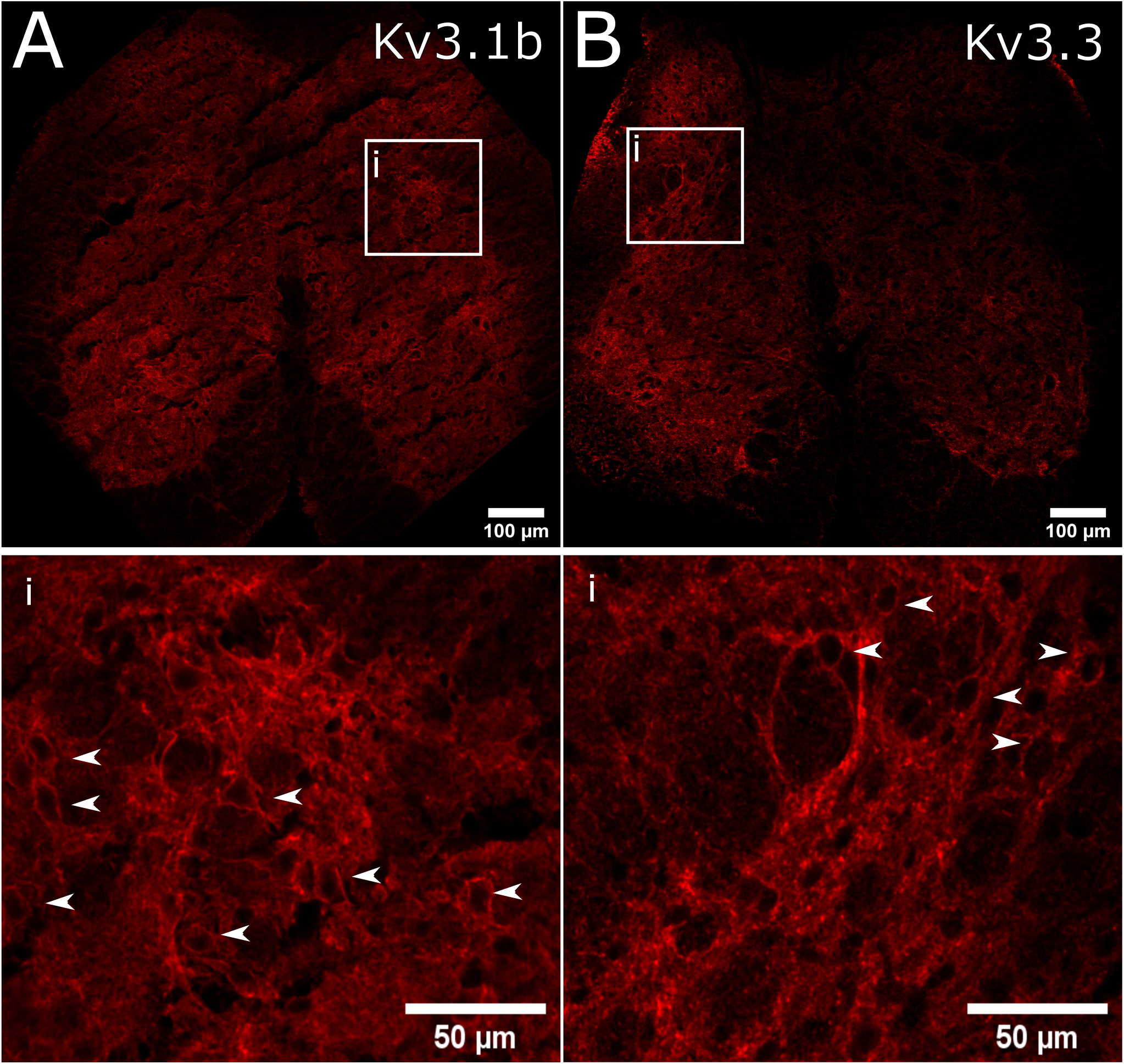
Immunofluorescent localisation of Kv3 subunits in intermediolateral lumbosacral spinal cord. **(A, B)** Immunofluorescence of Kv3.1b and Kv3.3 subunits. The white box outlines the intermediolateral autonomic area of the lumbosacral spinal cord and the right panel represents magnified images within this region. White arrows indicate Kv3 subunit immunofluorescence that appeared to encircle cells.

### Identification of a fast firing neuronal population, e-type 3, in the parasympathetic autonomic zone

Neurons in the parasympathetic autonomic zone lack a robust characterisation. We, therefore, performed an initial characterisation based on the firing phenotype to understand the different neuronal types and identify putative Kv3-positive fast firing cells in this area. Neurons were clustered according to a subset of electrophysiological characteristics such as firing frequency, adaptation index, inter spike interval coefficient of variation (synonymous with regularity), occurrence of delays, bursts, pauses and accommodation of firing (Fig 2A). Four distinct firing phenotypes, denoted e-types, were identified; e-type 1 steady regular, e-type 2 continuous adapting, e-type 3 fast bursting, e-type 4 steady delayed (Fig 2C). E-type 3 had a markedly higher rate of firing than the other e-types, which were indistinguishable based on firing rate alone (Fig 2B). E-type 3 had the briefest action potentials (1.3 ± 0.4 ms) and largest AHP amplitude (21.6 ± 5.9 mV), classical indications of a contribution of Kv3 current (Table 2). Indeed, k-means clustering of action potential waveforms for each cell revealed four AP-types. The briefest AP-type correlated most with the fast firing e-type 3 (46 %), however AP waveform clustering did not fully overlap with previous e-type clusters defined by firing properties, suggesting waveform shape alone cannot be used to identify each neuronal type (Supplementary1). In this initial characterisation, we identified a fast firing subpopulation and three slower neuronal phenotypes.

**Figure 2.**
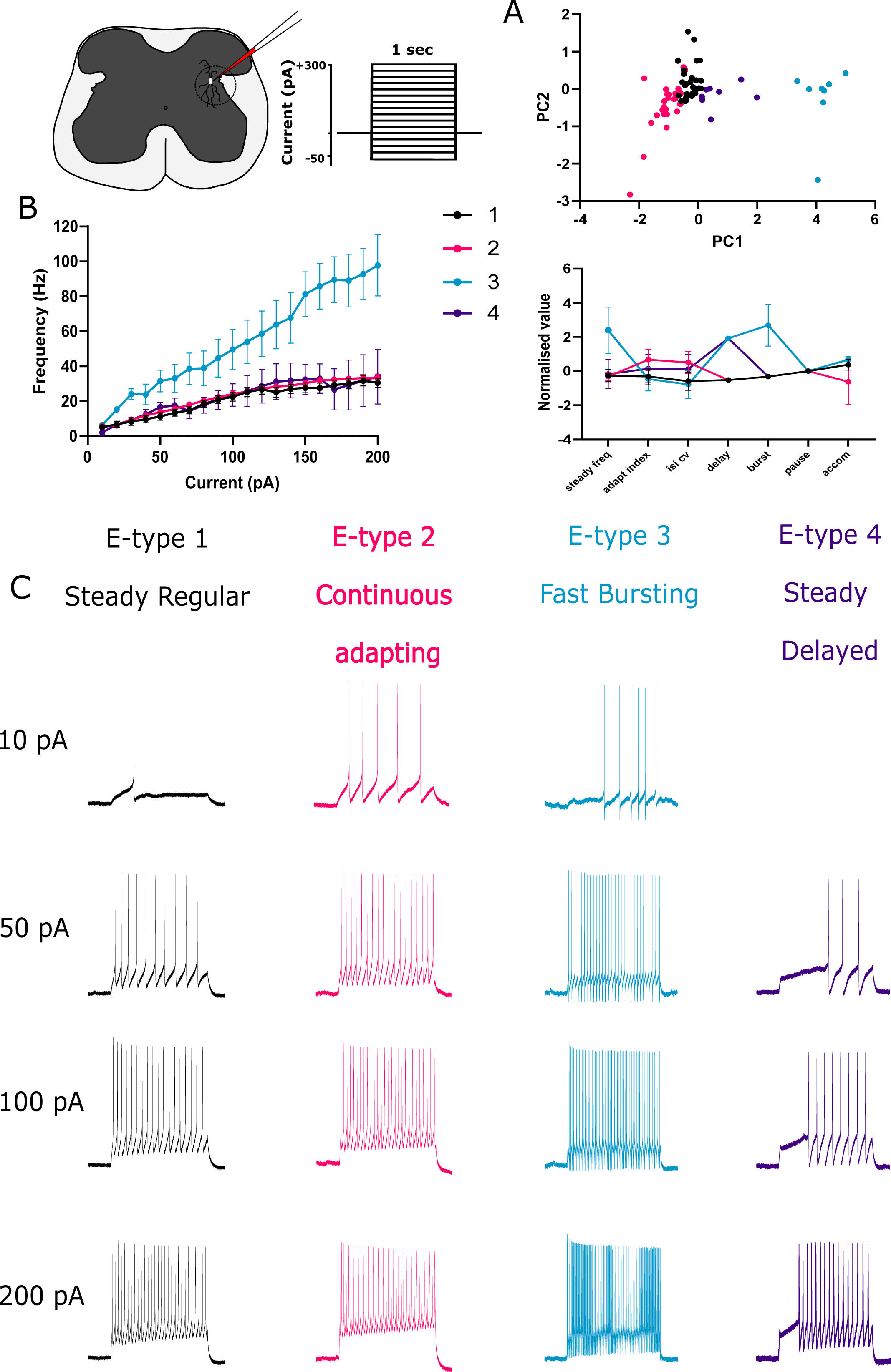
E-type classification of lumbosacral spinal neurons in lamina VII. **(A)** Neurons were clustered according to firing phenotype at maximal firing; steady state firing frequency, adaptation index, inter-spike interval coefficient of variation (ISI CV), delays, bursts, pauses and accommodation were used as clustering variables. Upper panel: Principle component scatter plot summarising clusters. Lower panel: Parallel plot comparison across clustering variables, error bars represent SEM. **(B)** Frequency-current plot summarising the firing frequency elicited by 1 second current steps from -70 mV for different e-types-note the fast firing e-type 3. **(C)** Four different e-types; steady regular, continuous adapting, fast bursting and steady delayed. Example traces for each e-type with 10, 50, 100, and 200 pA 1 second current injections. No firing phenotype was produced for e-type 4 at 10 pA.

**Table 2.** Description of action potential and afterhyperpolarisation shape

|  | **E1** | **E2** | **E3** | **E4** |
| --- | --- | --- | --- | --- |
| **AP Width (ms)** | $3.3 \pm 1.4$ | $4.0 \pm 1.8$ | $1.3 \pm 0.4$ | $3.5 \pm 2.0$ |
| **AP Amplitude (mV)** | $55.2 \pm 11.9$ | $52.0 \pm 13.9$ | $54.6 \pm 9.3$ | $45.4 \pm 10.4$ |
| **Repolarisation Duration (ms)** | $2.4 \pm 1.1$ | $2.9 \pm 1.2$ | $0.8 \pm 0.3$ | $2.4 \pm 1.6$ |
| **AP Threshold (mV)** | $-29.6 \pm 6.7$ | $-28.8 \pm 6.7$ | $-34.7 \pm 4.7$ | $-30.3 \pm 5.1$ |
| **AHP Duration (ms)** | $29.6 \pm 18.5$ | $28.8 \pm 16.3$ | $8.9 \pm 7.5$ | $27.1 \pm 20.6$ |
| **AHP Amplitude (mv)** | $19.5 \pm 5.3$ | $20.6 \pm 5.7$ | $21.6 \pm 5.9$ | $19.78 \pm 6.2$ |

### Morphological identification of PGN and interneurons

In order to differentiate between putative PGN and interneurons, where possible due to full morphological recovery, we assessed whether neuronal clusters had neurites projecting towards the ventral root, the site of efferent axonal exit from the spinal cord. This is an indicator of PGN (Morgan, 2002a). All e-types, except e-type 3, had neurons with clear ventral root projecting (VRP) neurites (Fig. 3A). This is further represented by plotting the average distance and angle (in 45 ° bins) of neurite end points from the soma for each e-type in the form of polar plots (Fig. 3C). E-types 1, 2, and 4 had long projections between 90 ° and 135 °, analogous to projecting towards the ventral root. Conversely, e-type 3 typically had a vertical orientation with short ventro-lateral projections and long dorsal projections. Kernel density estimates indicated the angular probability of projecting neurites; the peak angle for e-type 3 lay between 300 ° and 360 ° in the dorsal direction, whereas e-type 1 and 2 displayed prominent peaks in the lateral and dorso-medial direction, and e-type 4 had similar occurrences in all directions (Fig. 3C). E-type morphologies could not be significantly distinguished by Sholl parameters, ramification index (p = 0.13) and critical value (p=0.17, One-way ANOVA, Fig. 3B). However, this morphological assessment indicated that e-type 3 lacked a ventral-root projection and thus likely did not belong to an autonomic preganglionic motoneuron class.

**Figure 3.**
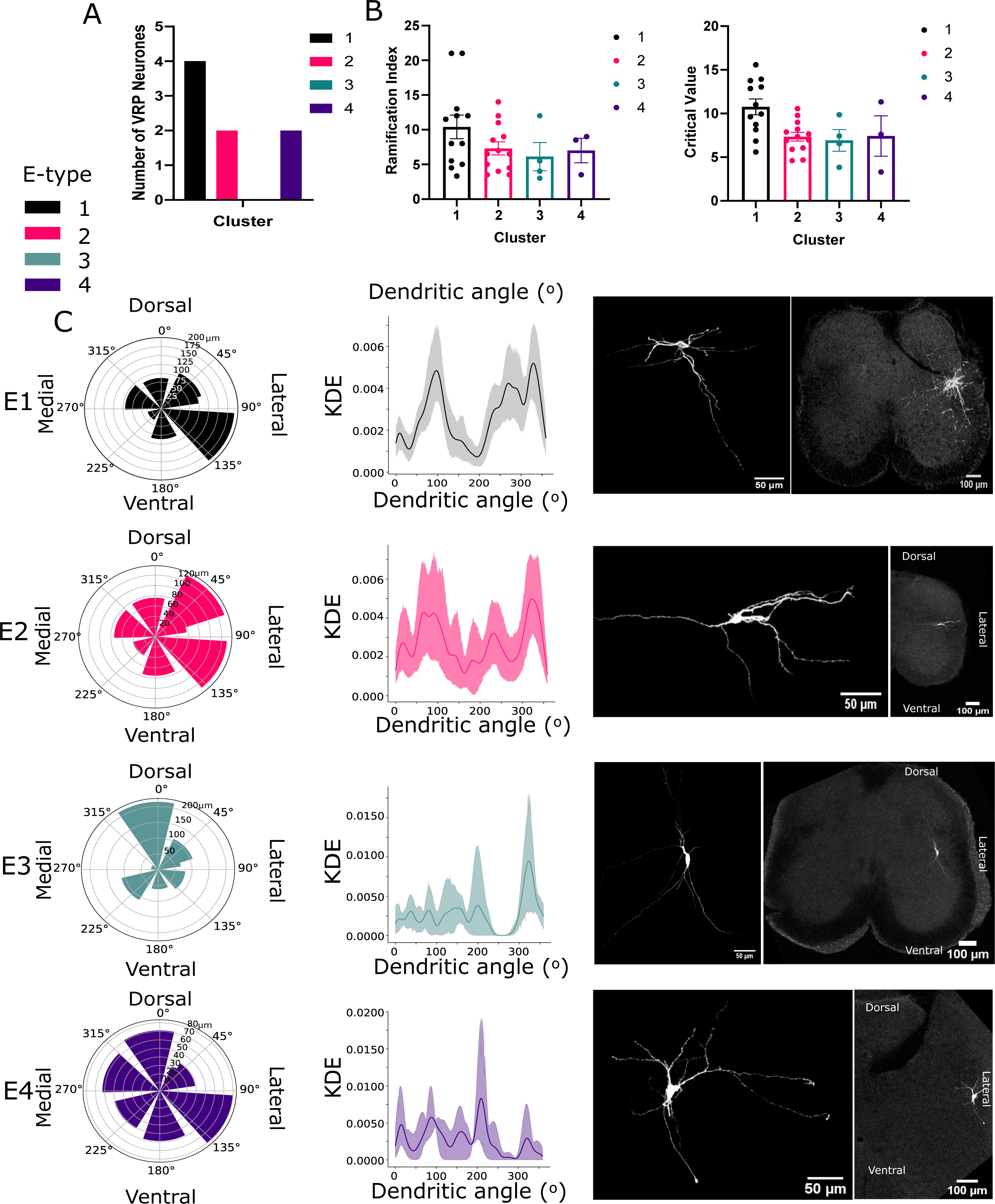
Distinct morphologies were associated with each e-type. **(A)** Comparison of the number of neurons with ventral root-projecting (VRP) neurites, typically associated with autonomic motoneurons. **(B)** Sholl analysis of reconstructed morphologies; ramification index and critical value between e-types. **(C)** Angles between the end of neurites and the soma were calculated to measure directional specificity (°) and spatial coverage (μm). Note most e-types had a large ventro-lateral projection except for e-type 3. Left panel: Polar plots indicating average dendritic length within 45 ° bins of dendritic end angle. Middle panel: Kernel density estimates indicating the directional likelihood of dendrites for each e-type. Right panel: Representative examples of reconstructed Neurobiotin-filled neurons for each e-type.

### Fast e-type 3 was able to follow high frequencies of stimuli

To test the responsiveness of each phenotype to slow and fast stimuli we injected 10 ms square current pulses at varying frequencies (20, 50 and 100 Hz) and amplitudes (160 -400 pA) and quantified the rate of failure to evoke an action potential for each pulse. Lower current amplitudes resulted in large failure rates as expected (Fig. 4A). E-type 3 had a significantly lower failure rate than the other e-types at 50 Hz and 100 Hz when compared at a high current amplitude (350 pA, mixed effects ANOVA, p = 0.0421, p=0.005, respectively). This indicated that e-type 3 was extremely responsive to high frequency stimulation (Fig. 4A-B), a strong suggestion of functional expression of Kv3 channels.

**Figure 4.**
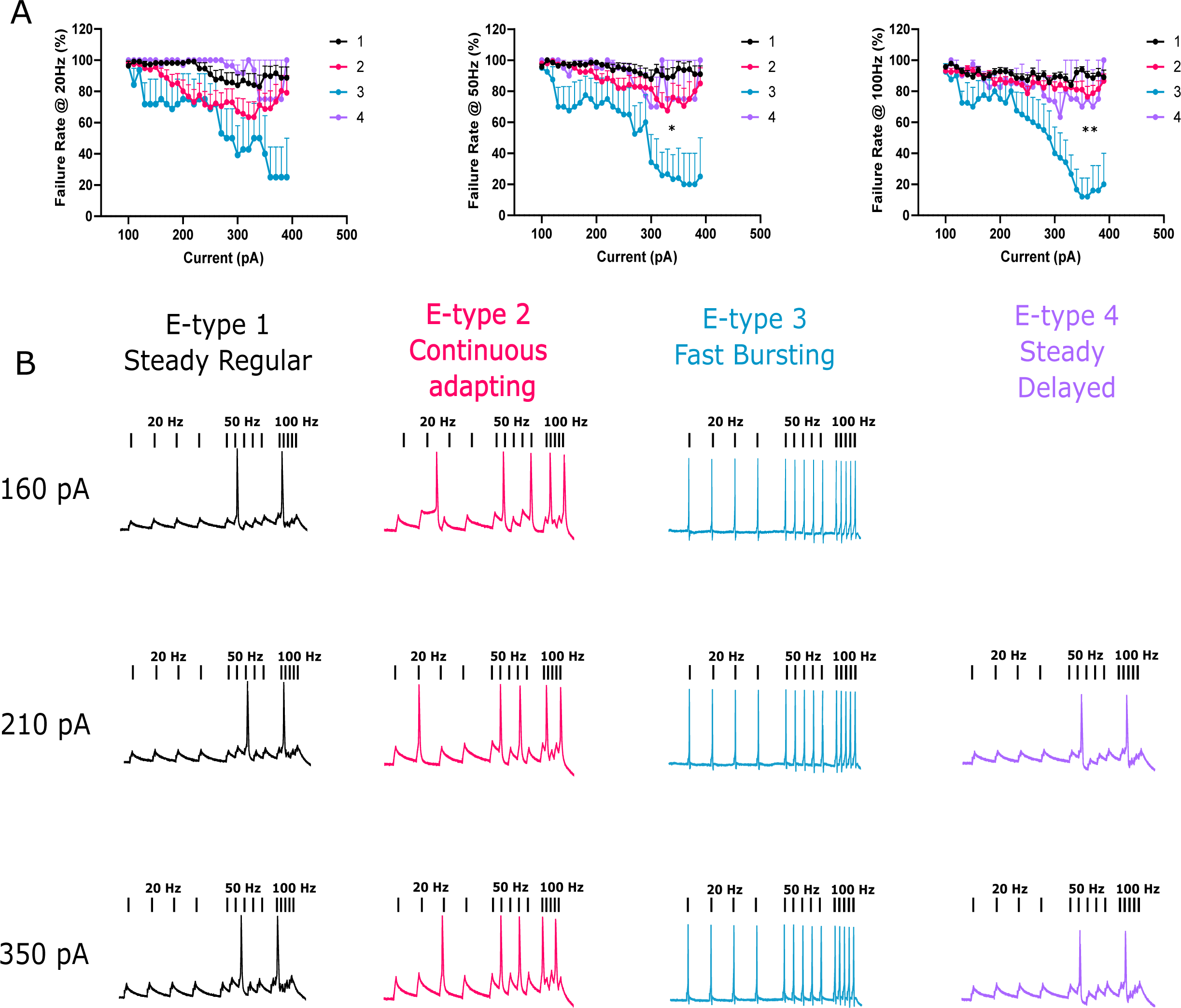
E-type 3 accurately followed high frequencies of stimulation. **(A)** 10 ms square current pulses were applied to neurons at different rates (20Hz, 50Hz and 100Hz) and at increasing amplitudes. Failure rate was quantified as a failure to fire an action potential during a pulse. E-type 3 successfully followed faster stimulation frequencies significantly better than other e-types. **(B)** Representative examples of action potentials evoked at different stimulation frequencies and at different current amplitudes for each e-type.

### Fast e-type 3 firing was impaired by application of TEA

We had uncovered a fast firing cell type, e-type 3, within the autonomic intermediolateral laminae. Furthermore, e-type 3 also displayed features prototypical of Kv3 channel expression, namely fast and responsive firing and brief action potentials. We next began to address whether Kv3 channels were crucial to these properties by using low concentrations of tetraethylammonium (TEA, 0.5 mM) to primarily block Kv3 channels (Fig. 5A, E1-4). Application of TEA significantly reduced firing frequency of e-type 3 by 24.5 Hz at 140 pA (multiple paired t-tests with Bonferroni correction, p = 0.0095, n=4, control; 60.25 ± 16.98 Hz, TEA; 35.75 ± 14.97 Hz ) (Fig. 5A E3).

**Figure 5.**
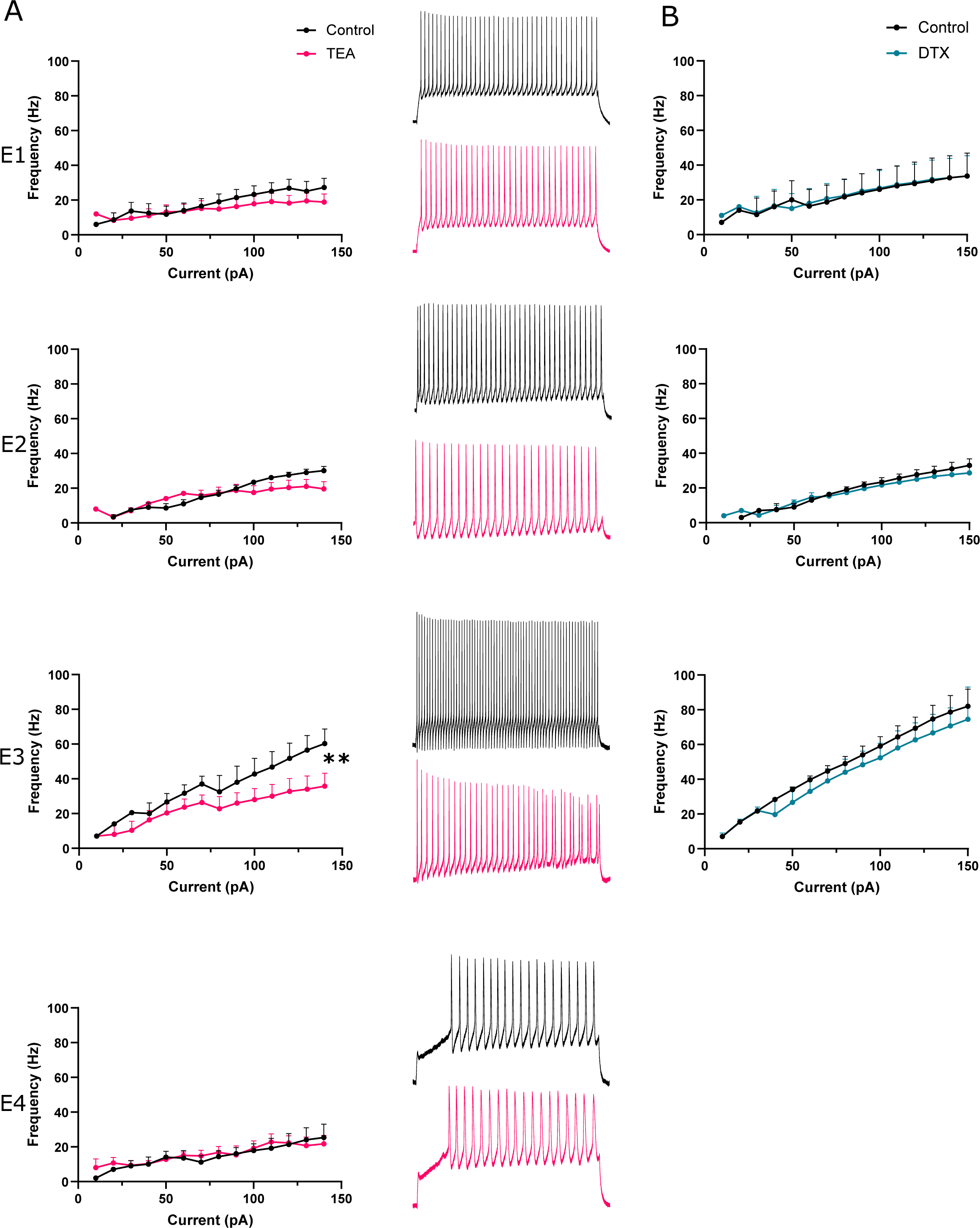
TEA impairs fast firing of e-type 3. **(A)** Left panel: Firing frequency-current plots for each e-type (1-4) in control aCSF and in presence of 0.5 mM TEA. Note significant reduction in firing rate at 140 pA in e-type 3. Right panel: Representative traces of neuronal firing in control aCSF and 0.5 mM TEA. **(B)** Firing frequency of each e-type over a range of current amplitudes in control aCSF and in presence of 10 nM DTX.

TEA (0.5 mM) had no significant effect on other e-types. TEA can also block Kv1 channels, however, application of Kv1 specific blocker dendrotoxin (DTX, 10 nM) had no discernible effect on firing frequency (Fig. 5B).

### A large proportion of e-type 3 outward current was TEA sensitive

Further interrogation with voltage clamp indicated low levels of DTX and TEA sensitive current across all e-types (Fig. 6A, B). Subtraction of currents evoked in 0.5 mM TEA from control was used to isolate the TEA-sensitive Kv3 component of total Kv current (Fig.6B). Interestingly, a large component of total Kv current was TEA and DTX insensitive, although this concentration of TEA does not produce a full block ( ∼ 50% ) of Kv3 channels (Coetzee et al., 1999; Rudy and Mcbain, 2001). TEA-sensitive current was significantly greater in e-type 3 at 14 mV compared to e-type 2 and 4 (One-way ANOVA, 3760 ± 1839 pA vs 955 ± 1012 pA and 991.7 ± 999.3 pA, p=0.0116 and p=0.04, respectively. Fig. 6A). TEA-sensitive current contributed a small proportion of total Kv current across other e-types. Activation of TEA-sensitive current for e-type 3 neurons was shifted by 18 mV towards more depolarised potentials compared to control (control V50 = -28.18 mV, TEA-sensitive V50 -10.66 mV, curve fit comparison p = 0.0012, extra sum of squares F test) suggesting that the TEA-sensitive current is produced by a high-voltage activated potassium channel, such as Kv3 channels. Here, through voltage clamp, we established that TEA-sensitive high-voltage activated putative Kv3 currents constituted a significant proportion of total Kv current in e-type 3 neurons and that it is most likely blockade of these channels with 0.5 mM TEA that reduced firing frequency.

**Figure 6.**
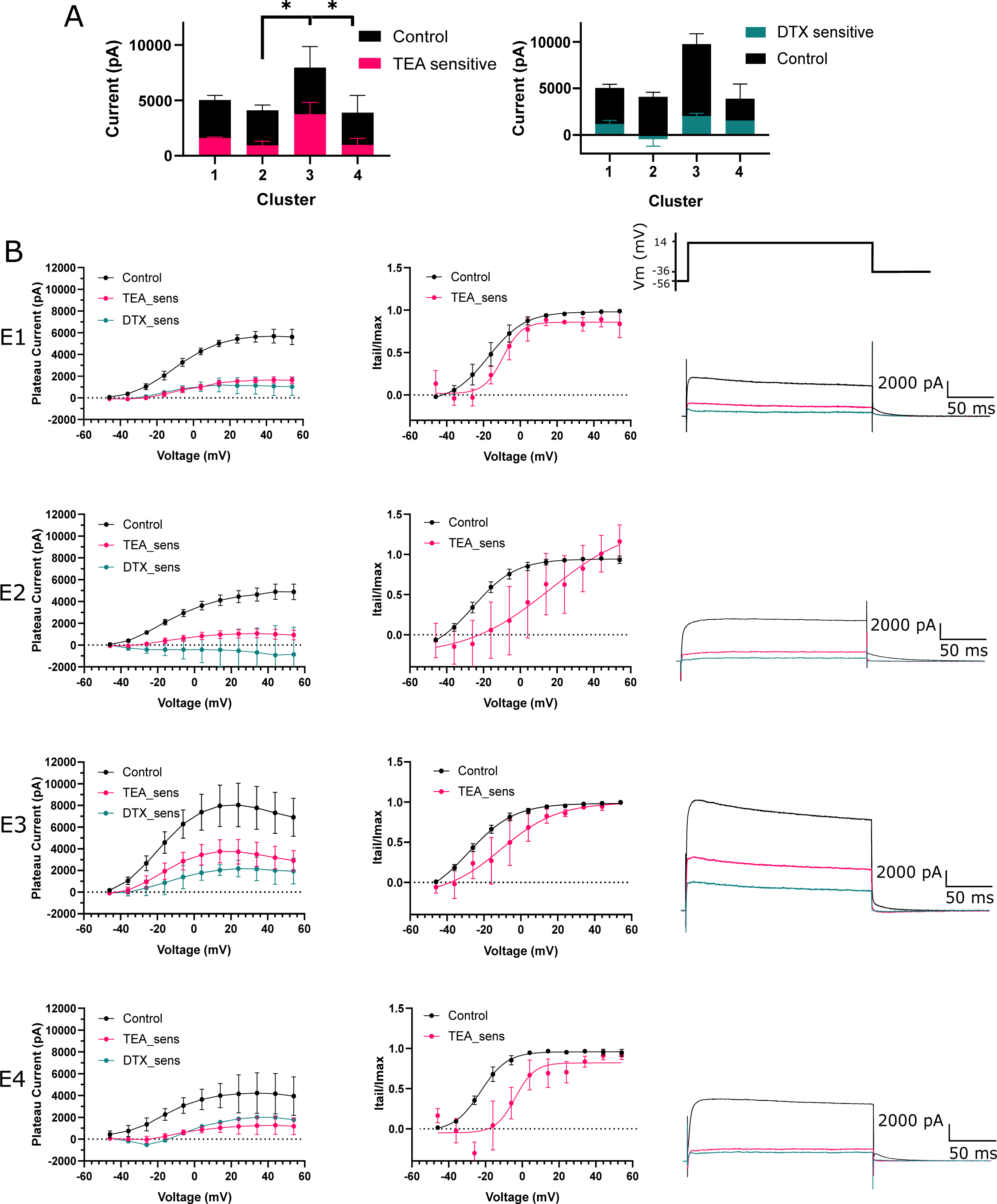
TEA sensitive Kv currents greatly contribute to outward currents in e-type 2. **(A)** Plateau outward currents evoked during a voltage step to 14 mV for each e-type in the presence of 0.5 mM TEA and 10 nM DTX. E-type 3 had significantly more TEA sensitive current than e-type 1. **(B)** Left panel: Voltage-current plots for each e-type (1-4). TEA and DTX sensitive (TEA_sens, DTX_sens) currents were obtained by subtracting currents in TEA and DTX from currents in control aCSF, the residual revealing the currents blocked by each compound. Middle panel: Activation plots indicating current activation over voltage steps. Current was measured during deactivating current tails and normalised to Imax. Right panel: Representative traces of outward currents evoked by a voltage step to 14 mV for each e-type. E-type 3 sustained a larger total current and TEA-sensitive current.

### Local and descending inhibitory input to PGN was TEA-sensitive

The data presented here provides strong evidence for the contribution of Kv3 channels to the rapid firing of a sub-population of neurons in the intermediolateral autonomic zone. However, Kv3 channels are also localised subcellularly and functionally important at the synaptic terminal, and likely to be important in any recurrent circuitry in the autonomic zone at the level of the synapse. To assess the role of Kv3 channels in synapses onto putative autonomic motoneurons in this region, such as e-types 1, 2 and 4, we evoked and recorded excitatory and inhibitory descending and local synaptic currents in the presence of TEA. 13 out of 15 neurons belonged to the putative autonomic neuronal classes. Descending fibres containing predomina<u>nt</u>tely excitatory (VGluT2) and some GABAergic and glycinergic descending bulbo-spinal input (Huma et al., 2014), were stimulated with a bipolar electrode placed in the lateral white matter of the lumbosacral spinal cord. Neurons were held at 0 mV to record IPSCs and at -56 mV to record EPSCs. Application of 0.5 mM TEA increased evoked IPSC (eIPSC) amplitude by more than 100% (Paired t-test p=0.03, control 94.93 ± 115.1 pA TEA 210 ± 134 pA, n=9. Fig. 7B). Paired-pulse stimulation was used to assess potential presynaptic site of action and TEA reduced the paired-pulse ratio by 33 % (p=0.0286, n = 6, Control 1.564 ± 0.59, TEA 1.07 ± 0.42, Paired T-test). Kv1 channels, also blocked by millimolar concentrations of TEA, are important regulators of synaptic excitability. To rule out a contribution to the observed effect we repeated the experiment in the presence of DTX (10 nM). DTX did not have a significant effect on eIPSC amplitude suggesting eIPSC potentiation was solely due to Kv3 blockade (Fig. 7B). Super-resolution Airyscan co-immunolabelling of GlyT2 and VGAT confirmed the presence of inhibitory (glycinergic and GABAergic) synaptic terminal markers in the intermediolateral laminae of the lumbosacral spinal cord and revealed a degree of overlap with Kv3.1b, (14% and 13 %, respectively, of boutons positive for Kv3.1b) (Fig 7C). We delivered a train of stimulus to further demonstrate TEA sensitive potentiation of eIPSC amplitude during continuous activity. Histogram analysis of a 10 Hz train of eIPSCs demonstrated a significant shift towards larger amplitude eIPSCs in the presence of TEA (Two sample Kolmogorov-Smirnov Test, p= 0.0018, Fig. 7D left). Further analysis of spontaneous background IPSCs (sIPSC), likely due to local activity, also produced a similar shift towards larger sIPSC amplitudes during application of TEA (Two sample Kolmogorov-Smirnov Test, p= 0.0004, Fig 7D right). Most synaptic responses depressed over the course of the 10 Hz train with a mean linear fit gradient of -2.66 ± 5.6. Lack of activity-dependent facilitation here suggested that the fast-inactivating Kv3 subunit Kv3.4, was not functionally expressed at these synapses and were not the target of the effect of TEA. Together, this data strongly indicated that Kv3 channels were expressed in inhibitory terminals and functionally constrained both descending and local inhibitory synaptic amplitude.

**Figure 7.**
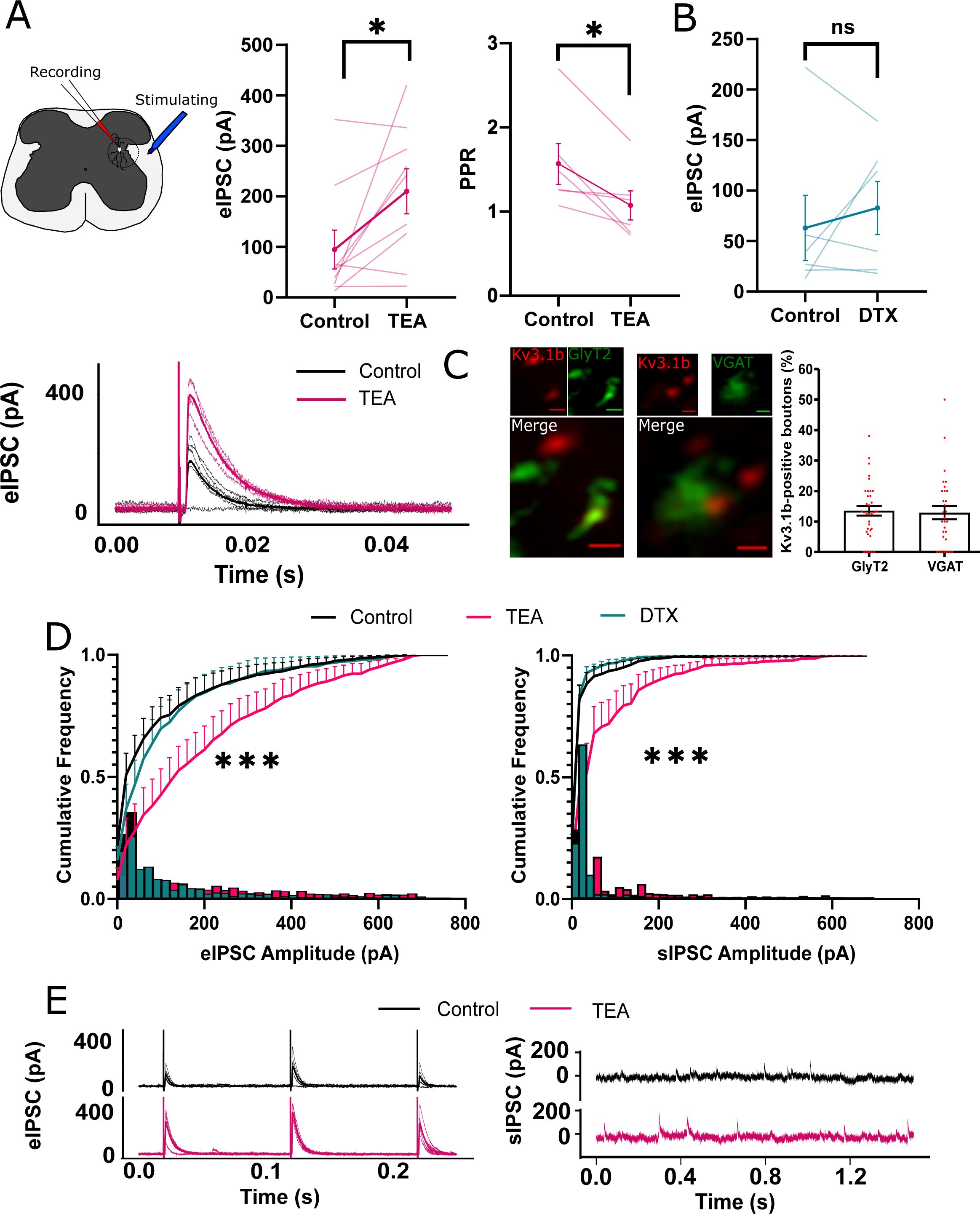
Kv3 blockade potentiates descending and local inhibitory synaptic responses. A bipolar electrode positioned in the lateral white matter was used to evoke descending synaptic currents. Neurons were held at 0 mV to isolate inhibitory post-synaptic currents (IPSCs). **(A)** Left panel: Evoked IPSC (eIPSC) amplitude between control and 0.5 mM TEA (each line represents an individual cell). Right panel: Paired pulse ratio between IPSCs 100 ms apart in 0.5 mM TEA. Bottom panel: Representative example of potentiated IPSC amplitude in TEA. **(B)** Effect of bath application of 10 nM DTX on eIPSC amplitude. **(C)** Left panel: super resolution Airyscan images showing clear overlap of example synaptic boutons with Kv3.1b IF with GlyT2 IF. Kv3.1b co-localisation with inhibitory markers GlyT2 and VGAT represented as the number of Kv3.1b positive boutons. **(D)** Cumulative (line) and relative frequency (bar) plots of eIPSC (left panel) and spontaneous background IPSC (sIPSC) amplitudes in the presence of 0.5 mM TEA and 10 nM DTX. **(E)** Representative traces of eIPSCs (left panel) and sIPSCs (right panel) in control and 0.5 mM TEA.

### Local and descending excitatory input to PGN was TEA-insensitive

Conversely, application of TEA had no significant effect on eEPSC amplitude and PPR (Fig. 8A). Interestingly, incubation in DTX significantly decreased eEPSC amplitude (control; 134.2 ± 72.9 pA, DTX; 92.78 ± 53.90 pA, paired t-test, p = 0.047, Fig. 7B) and shifted cumulative frequency EPSC amplitude towards smaller amplitudes for evoked but not spontaneous EPSCs (Fig. 8C, Two sample Kolmogorov-Smirnov Test, p= 0.001). This implied that blocking Kv1 channels decreased synaptic amplitude only in descending excitatory input. Immuno-labelling revealed a degree of overlap between Kv3.1b and VGluT2 immunoreactivity (12%) (Fig 8E) suggesting expression of Kv3 channels in a small proportion of excitatory synaptic terminals in this area. The combined analyses presented here suggests that Kv3 channels are expressed in the somatic and synaptic membrane of inhibitory interneurons in the intermediolateral autonomic zone of the lumbosacral spinal cord, and that here they facilitate a distinctive fast firing phenotype and constrain inhibitory input to putative PGN. Consequently, Kv3 channels may represent an important identifier of fast polysynaptic or recurrent inhibition of PGN.

**Figure 8.**
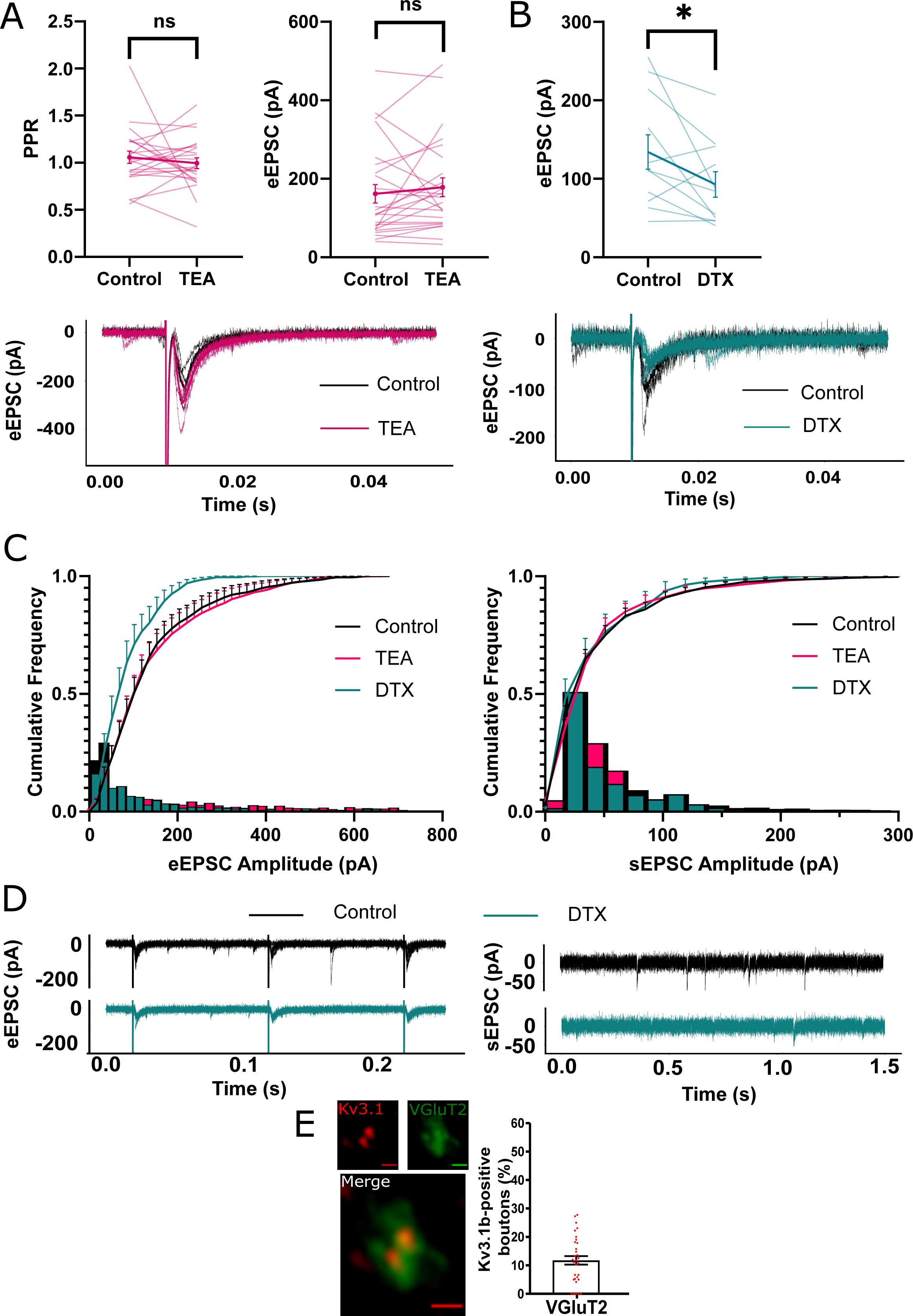
Kv3 blockade has no effect on excitatory synaptic responses. Neurons were held at -56 mV to isolate excitatory post-synaptic currents (EPSCs). **(A)** Left panel: Evoked EPSC (eEPSC) amplitude between control and 0.5 mM TEA (each line represents an individual cell). Right panel: Paired pulse ratio between EPSCs 100 ms apart in 0.5 mM TEA. Bottom panel: Representative example of unaffected EPSC amplitude in TEA. **(B)** Upper panel: Effect of bath application of 10 nM DTX on eEPSC amplitude. Lower panel: Representative example of eEPSC in control and DTX. **(C)** Cumulative (line) and relative frequency (bar) plots of eEPSC (left panel) and spontaneous background EPSC (sEPSC) amplitudes in the presence of 0.5 mM TEA and 10 nM DTX. **(D)** Representative traces of eEPSCs (left panel) and sEPSCs (right panel) in control and 10 nM DTX. **(E)** Example Airyscan images and bar chart showing limited co-localisation of VGluT2 boutons with Kv3.1b.

## Discussion

### The identity of e-types

We identified four neuronal sub-populations in the lumbosacral intermediolateral spinal cord based on electrophysiological criteria; one type, e-type 3, represented a fast-firing population and the remaining e-types, with longer AP duration and slower firing properties most likely represented parasympathetic motoneurons. We postulate that e-type 3 represented an interneuronal population for two main reasons; 1) we never observed a ventral root projecting neurite, typically indicative of PGN (Morgan, 2002a), and 2) e-type 3 neurons fired faster than reported frequencies for PGN (de Groat and Ryall, 1968a). Consequently, according to these criteria, e-types 1, 2 and 4 likely represent autonomic subpopulations of which multiple sub-types have indeed been postulated (Morgan et al., 1993). This is an important characterisation that will improve our understanding of the parasympathetic function of the spinal cord, however it is important to note that the machine learning method of clustering used here is far from deterministic and represents a best estimate at the number and defining characteristics of subpopulations. However, by beginning to name specific neuronal types in this autonomic region, as has been accomplished in the dorsal horn (Todd, 2017), we can begin to ask what role they play in crucial physiological processes that involve parasympathetic input from the spinal cord, as is seen during the micturition reflex.

### Kv3 channels are crucial for the electrophysiological characteristics of e-type 3

We further examined the biophysical underpinnings of the phenotype of e-type 3 with the hypothesis that expression of Kv3 channels underlie the fast firing ability. Kv3 channels are well known to confer rapid firing properties in multiple other CNS regions (Erisir et al., 1999; Lien and Jonas, 2003). Furthermore, association of the Kv3.1b subunit with intermediolateral fast firing interneurons in the thoracic spinal cord has been previously established (Deuchars et al., 2001; Brooke et al., 2002). In this study, application of Kv3 channel blocker, TEA, reduced firing frequency and isolated high-voltage activated outward currents that constituted a large proportion of total outward current. Furthermore, neuronal expression of Kv3 subunits in this area confirmed the importance of Kv3 channels in the physiology of this neuronal class. However, functionally distinguishing contributions of Kv3 subunits to the firing phenotype of e-type 3 is difficult without selective inhibitors for each subunit. This would require the use of knockout Kv3 mouse lines in future experiments to evaluate the contribution of each subunit to the phenotype of e-type 3 as well as the effect on bladder control (Choudhury et al., 2020). Kv3 channels have been selectively associated with inhibitory populations in ventral and dorsal regions; Kv3 subunits are expressed in neurons that fire at high frequency such as Renshaw cells that inhibit motoneuron output (Song et al., 2006) and inhibitory dorsal horn interneurons that gate sensory input (Nowak et al., 2011). Are Kv3 channels also associated with inhibitory populations in the intermediolateral autonomic zone? Viral tracing studies show that interneurons synaptically connected to PGN are in the autonomic zone and focal extracellular stimulation of these local interneurons evoke both EPSCs and IPSCs in PGN (Araki, 1994; Araki and De Groat, 1996; Nadelhaft et al., 2000; de Groat et al., 2015). However, the TEA-sensitivity of local inhibitory but not excitatory inputs onto putative PGN suggests that Kv3 channels are associated with inhibitory populations synaptically connected to PGN. Our data did not distinguish an average difference in local and descending excitatory input amplitude after Kv3 blockade, but some examples are identifiable within out dataset that show potentiated excitatory synaptic responses. Further, Kv3.4 subunits have been associated with sensory glutamatergic afferent terminals in the dorsal horn (Muqeem et al., 2018), thus, we cannot completely rule out association of Kv3 channels with excitatory neurons. Indeed, our attempts to address this question directly using paired recordings within the intermediolateral region found limited connectivity, thus we could not distinguish between excitatory and inhibitory populations synaptically connected to PGN. The role of Kv3 channels at the synaptic terminal is well established, where application of Kv3 channel blockers broadens the presynaptic action potential, increases Ca^2+^ influx and neurotransmitter release (Ishikawa et al., 2003; Goldberg et al., 2005; Kuznetsov et al., 2012; Rowan et al., 2014). Our findings further corroborate this role in the spinal cord, however, with a degree of selectivity for inhibitory terminals indicating that Kv3 channels are more likely to constrain inhibitory output. The punctate immunofluorescence of Kv3.1b and Kv3.3 suggests both subunits may be important in presynaptic dynamics. We observed no activity dependent facilitation of post synaptic currents, but whether the inactivation of Kv3.3 is insufficient to elicit such a mechanism, as has been demonstrated for Kv3.4, is not known (Rowan and Christie, 2017).

### Other distinct ionic contributions to e-types

We primarily focussed on Kv3 channels in fast firing neurons, however there are clear indications of other ionic currents within our data. All e-types displayed a prominent voltage deflection “sag” characteristic of Ih during hyperpolarising current injections, likely mediated by HCN channels. The delayed firing of e-type 4 is highly suggestive of IA typically conducted via Kv4 channels. Indeed, other autonomic neurons such as sympathetic preganglionic neurons have been found to express IA, where it endows low-pass filtering of incoming excitatory synaptic input by decreasing EPSP summation between 15 and 40 Hz, thus reducing transmission of higher frequency activity (Briant et al., 2014). This was reflected in e-type 4 being unable to follow higher frequencies of stimuli and thus a similar low pass filter may exist in PGN. Voltage clamp data from all e-types also indicated a high-voltage activating delayed rectifier with slow activation that was TEA and DTX-insensitive often indicative of expression of Kv2 channels (Johnston et al., 2010). The slow kinetics of Kv2 channels ensure AP amplitudes are consistent by ensuring hyperpolarised inter-spike potentials and thus a stable pool of available non-inactivated NaV channels (Johnston et al., 2008). This ubiquitous expression likely explains the consistent AP amplitudes observed in all our e-types.

### An autonomic Renshaw cell?

A fundamental question is what is the purpose for an inhibitory neuronal class to fire so much faster than its autonomic neighbours? A possible hypothesis is to enable rapid recurrent inhibition of PGNs to suppress activation of the parasympathetic motoneuronal pool and thus suppress contraction of the bladder detrusor smooth muscle. PGN extend axon collaterals within the spinal cord indicating a likely recurrent activation of other spinal neurons (Felkins and Zhang, 1991; Morgan, 2002b). Furthermore, a recurrent inhibitory reflex akin to that of the motoneuron-Renshaw cell motif has been described for PGNs (de Groat and Ryall, 1968b). This inhibition was shown to be glycinergic in nature, was mediated by fast firing interneurons in the immediate vicinity of PGNs and depressed bladder detrusor contractions. The similar localisation and phenotype of the e-type 3 class suggests that this neuronal population could represent a candidate “autonomic Renshaw cell”. Indeed, previous research using rabies virus identified Kv3-expressing neurons antecedent to autonomic neurons in the thoracic spinal cord (Deuchars et al., 2001; Brooke et al., 2002). Furthermore, Kv3 channels consistently correlate with the fast firing inhibitory element of recurrent inhibitory circuits, from in Renshaw cells to cortical neurons (Song et al., 2006; Espinosa et al., 2008). Renshaw cells are thought to limit or terminate the activity of somatic motoneurons as well as potentially determine selective excitability of motoneurons (Noga et al., 1987). Analogously, an autonomic Renshaw cell, could completely suppress PGN activity and bladder contraction. However, this doesn’t seem to be the case when the volume of the bladder is high (de Groat and Ryall, 1968b) but perhaps represents a mechanism of PGN quiescence during bladder filling and continence. Alternatively, with the expression of Ih current in putative PGN e-types, recurrent inhibition could define a rhythmic pattern of activity through rebound depolarisation (van Hook and Berson, 2010), a pattern that could contribute to the spontaneous contractions observed for bladder detrusor muscle. Additionally, recurrent inhibition could allow selective activation of PGN with the greatest degree of excitation within the autonomic pool, where activity of the most excited PGN suppresses weakly excited PGN to better coordinate organs of the lower urinary tract during bladder voiding. These hypotheses assume multiple strong inhibitory inputs proximal to the soma to suppress PGN firing, however distal inputs may instead shunt dendritic afferent input as has been observed for Renshaw cells and motoneuron dendrites (Bhumbra et al., 2014). Kv3 channels have become pharmacological targets for a variety of neurological disorders in the past decade, with novel compounds typically acting to slow the rate of firing of Kv3 expressing neurons or recover the firing rate in impaired paradigms (Rosato-Siri et al., 2015; Parekh et al., 2017; Brown et al., 2020). A pharmacologically-induced reduction in the rate of inhibition to PGN could confer increased parasympathetic activity and unwanted bladder detrusor contractions during bladder filling and typical periods of continence. However, there is some evidence of reductions of Kv3 subunits in the CNS during ageing, and these may lead to impaired function of fast firing neurons (Zettel et al., 2007). Perhaps firing-impaired interneurons in the spinal cord could disrupt spinal parasympathetic dynamics that correlate with age-related decline in bladder control and be amenable to pharmacological recovery. By fundamentally characterising the individual neuron types within this circuitry and the biophysical mechanisms that underpin the phenotype of those cells, we can begin to understand and modulate spinal parasympathetic dynamics during important physiological reflexes such as the micturition reflex.

## Acknowledgements

Acknowledgements

This work was supported by BBSRC grant BB/L01565X/1. Pierce Mullen’s current affiliation is with School of Psychology and Neuroscience, University of St Andrews.

---

**Figure S1.**
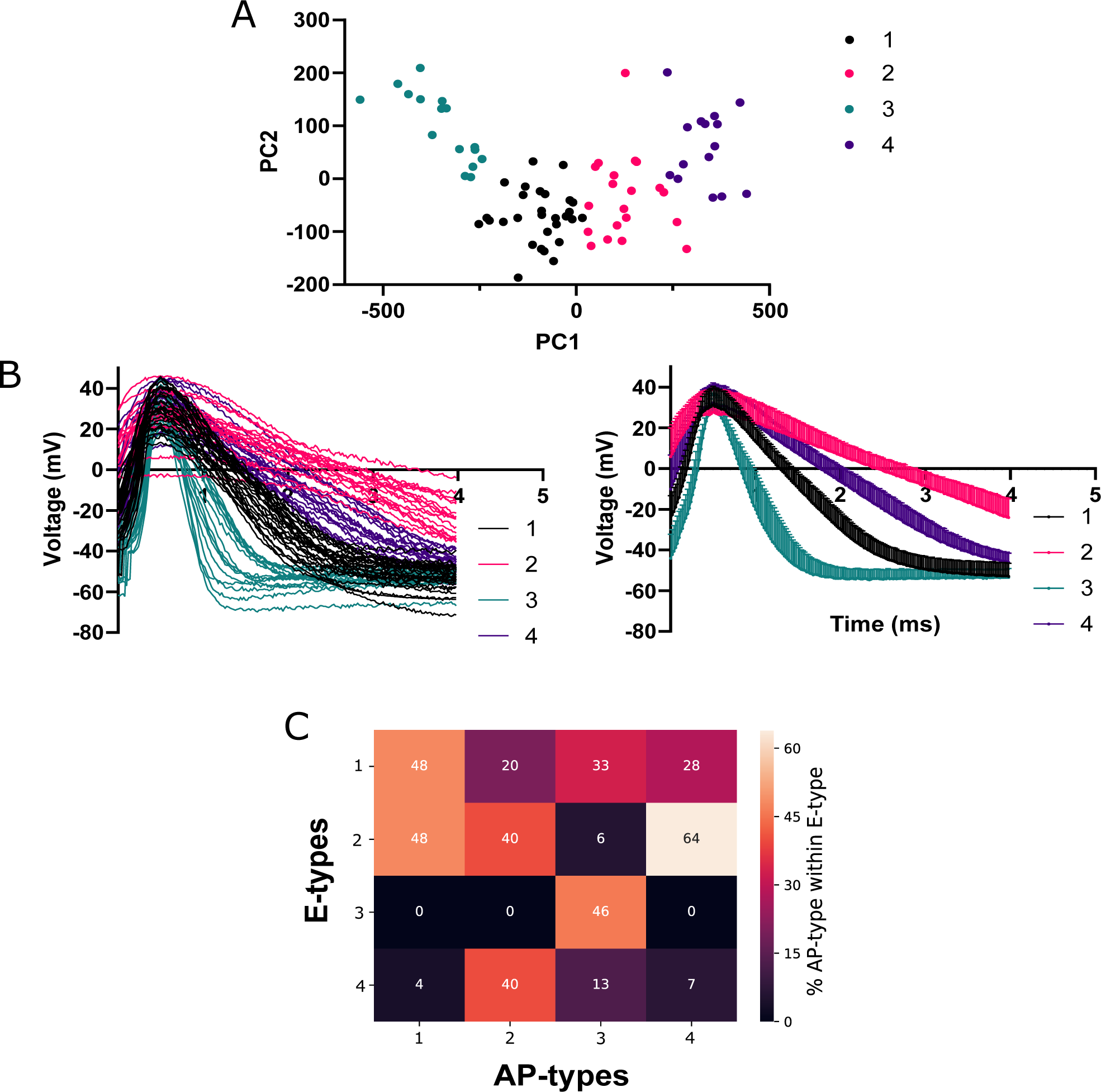
Correlation of AP waveform with e-types. **(A)** The first action potential waveform produced at rheobase for each neuron was clustered. Scatterplot of principal components 1 and 2 indicating summarising clusters. **(B)** Individual (left panel) and average waveforms (right panel) for each spike type. **(C)** Correlation of AP spike type with e-types, represented as the percentage of AP-types associated with each e-type.

